# Different underlying mechanisms for high and low arousal in probabilistic learning in humans

**DOI:** 10.1101/2021.02.26.431959

**Authors:** Luis F. Ciria, Marta Suárez-Pinilla, Alex G. Williams, Sridhar R. Jagannathan, Daniel Sanabria, Tristán A. Bekinschtein

## Abstract

Humans are uniquely capable of adapting to highly changing environments by updating relevant information and adjusting ongoing behaviour accordingly. Here we show how this ability —termed cognitive flexibility— is differentially modulated by high and low arousal fluctuations. We implemented a probabilistic reversal learning paradigm in healthy participants as they transitioned towards sleep or physical extenuation. The results revealed, in line with our pre-registered hypotheses, that low arousal leads to diminished behavioural performance through increased decision volatility, while performance decline under high arousal was attributed to increased perseverative behaviour. These findings provide evidence for distinct patterns of maladaptive decision-making on each side of the arousal inverted u-shaped curve, differentially affecting participants’ ability to generate stable evidence-based strategies, and introduces wake-sleep and physical exercise transitions as complementary experimental models for investigating neural and cognitive dynamics.

## INTRODUCTION

Making mistakes is inherent to learning and the accomplishment of any task. We make mistakes every day, even when faced with the same task repeatedly. Our ability to learn from these errors and flexibly adapt ongoing behaviour according to changes in the environment is critical for our survival. This ability —termed cognitive flexibility— depends on our innate capacity to establish associations between stimuli (S), responses (R), and outcomes (O), as well as to integrate previously acquired knowledge and skills into effective strategies for coping with similar future demands.^1^ Here, we implement a Probabilistic Reversal Learning (PRL) task to study the modulatory effect of low and high arousal on cognitive flexibility — participants continue to perform as they fall asleep or with increasing physical exercise— to map either side of the Yerkes-Dodson Curve (1908).^2^

Cognitive flexibility is often studied using PRL tasks, typically assigning probabilistic reinforcement contingencies to abstract S-R associations, that are later abruptly reversed, requiring participants to learn new S-R reinforcement contingencies by trial and error to overcome prepotent ones^3^. Efficient performance relies on learning from the reinforcement received^4^, the estimation of the likelihood that a reversal may occur,^5,6^ and the continuous integration of a history of choices and reinforcements.^7^ Indeed, evidence from both human and animal studies suggests that different high- and low-order strategies or series of rules are adopted during reversal learning, leading to maladaptive response patterns when the external pressures change or when the internal milieu varies.^7,8^ Parsing the microstructure of learning derived from trial-by-trial responses enables the dissociation of the cognitive processes and behavioural strategies that drive subjects’ choices during reversal learning. Here we propose that arousal fluctuations may differentially modulate cognitive flexibility leading to distinct maladaptive behavioural patterns of performance.^9^

Fluctuations in arousal and alertness (hereafter described jointly as “arousal”) occur constantly across the day but are exacerbated during transitions toward strained states such as sleep^10^ or physical extenuation,^11^ where arousal levels change drastically in a progressive and nonlinear manner.^12,13^ These arousal fluctuations play a crucial role in modulating cognition, facilitating or hindering certain cognitive processes and performance to internal and external stimuli. ^14,15,16,17,18^

The interaction between arousal and cognition has been traditionally approached from the perspective proposed by Yerkes and Dodson in 1908.^2^ According to their famous inverted U-shaped law, the optimal level of cognitive performance in complex tasks is reached at moderate levels of arousal, whereas deviations from this optimal arousal point, below or beyond, result in cognitive performance impairments. Though reductionist, the inverted U-shaped law represents a useful minimal framework to characterize the neural and cognitive dynamics of many physiological states across the arousal spectrum. Among these physiological states, researchers have paid special attention to reduced arousal states, including sleep stages,^19^ sedation,^20^ sleep deprivation,^21^ motivation^22^ and fatigue.^23^

Sleep can be used as the gold standard model of transition toward low arousal.^10^ This area looking at the interaction between homeostasis and cognitive function is understudied due to the complexity of capturing dynamically metastable states like mild sedation^24,25^ and drowsiness.^17^ When falling asleep, individuals manifest a wide range of changes, from physiological to phenomenological, that are categorized into several well-described sleep stages.^26^ One of these stages is drowsiness, a transitional stage of consciousness between attentive wakefulness and light sleep, characterized by a progressive and nonlinear loss of responsiveness to external stimuli which does not immediately imply unconsciousness.,^27,28,29^ Drowsiness, as well as similar reduced arousal states, has been repeatedly associated with an impairment of cognitive processing, and particularly the capacity to deal with conflicting information,^18^ attentional performance,^30^ and perceptual decision-making.^31^ However, in drowsiness, and even during highly reduced arousal states, pre-attentive and early bottom-up attentive processing can still be accomplished with and without conscious awareness. ^17,32,33^

The transition towards the other side of the arousal spectrum (i.e., heightened arousal states) has received even less attention.^34^ The absence of a theoretical model for progressive physiological transitions towards high arousal states, has also contributed to a lack of advance in the field. Here, we consider endurance physical exercise as a useful experimental model of arousal transition upwards, with many commonalities with sleep transition. A single bout of endurance physical exercise (e.g., running or cycling) up to physical extenuation involves a complex transition encompassing a wide range of changes (e.g., neural, motor, endocrinal, phenomenological, etc.), that are also categorized into several well-described stages, from resting, through the aerobic and the anaerobic thresholds, up to the limit where the individual has to stop.^35^ This highly fluctuating transition has been also associated with changes in cognitive processing to internal and external stimuli.^36,37,38^ In particular, high-order top-down processes that govern goal-directed behaviour in changing environments (i.e., cognitive control) appear to benefit from increases in the level of arousal^39^ up to a certain exercise intensity. Further intensity increments approaching and exceeding the anaerobic threshold seem to hinder cognitive performance,^36,37,38,40^ in line with the Yerkes-Dodson law prediction.

Sleep and physical exercise provide complementary perspectives on the cognitive dynamics, and experimental models, when the arousal level is altered. However, and despite the fact that both sides of the arousal spectrum exhibit similar cognitive performance impairments, they cannot be treated as mirroring states in terms of cognitive performance without a fine-grained differentiation of the behavioural dynamics that lead to these global impairments. Furthermore, the theoretical differences in the transitions towards sleep or complete (physical) exhaustion have to be considered in the assumptions and interpretations of this and future studies. Thus, it is crucial to ask when arousal is altered (increased or decreased), which specific processes of cognitive flexibility and information processing are affected, and whether low and high arousal states are characterized by different strategic behaviours underlying decision-making. It should be understood that the physiological processes underlying the change in performance seen in different Dodson-Yerkes experiments since 1908 are different at each side of the curve, and it should be expected that these changes in arousal modulate differently the cognitive abilities. Here, we use a PRL task to disentangle the behavioural dynamics of cognitive flexibility as they get modulated by ongoing fluctuations in arousal levels and to further delineate the microstructure of learning derived from trial-by-trial responses to conflicting evidence. In particular, we manipulated arousal level to facilitate natural transitions to low alertness, from awake to asleep; or to elicit high arousal, instructing participants to exercise during 60 minutes at the highest intensity and effort possible without reaching premature exhaustion. During both arousal modulations, participants performed a PRL task, requiring the adaptation of behaviour following changes in reinforcement and punishment, as well as the maintenance of strategic response patterns in the face of misleading (probabilistic) feedback.

Based on the premises that (1) drowsiness hinders the extraction of task-relevant information from external stimuli and its integration, fragmenting specific aspects of cognition while preserving crucial executive control processes;^18,31,33,41^ (2) drowsiness has been associated with more liberal decision-making;^17,30,31^ (3) moderate-to-high intensity endurance exercise leads to a selective enhancement of executive control processes while lower and higher intensities result in an impairment or minimal effect;^40,42,43^ and (4) high arousal promotes habitual responding and reduced engagement of complex cognitive strategies;^44,45,46^ predicted that behavioural performance would be enhanced in moderate-intensity physical exercise, while drowsiness and high-intensity exercise would lead to diminished performance in light of the inverted U-shaped Yerkes-Dodson Law. Specifically, we hypothesized that reduced arousal states would be associated with an impairment of performance (compared to baseline), which would be attributed to a tendency to apply a simple strategy (win-stay/lose-shift) instead of using an integrated history of choices and outcomes to drive performance (probabilistic switching behaviour). In contrast, while we also expected an impairment of performance during heightened arousal states, we hypothesized it would be attributed to a failure to disengage from ongoing behaviour (perseveration). In addition, we hypothesized that altered arousal states might reduce the ability of participants to apply a proper higher order strategy, resulting in wide periods of time-on-task in which participants would perform the task simply responding to the tones (i.e., automatic rule) but without applying any strategy (i.e., higher order rule). All these hypotheses, together with the analysis plan, were pre-registered after data collection.^9^

## RESULTS

To investigate the modulatory effect of arousal fluctuations on cognitive flexibility, a PRL task was carried out with human participants (n=100) while they were transitioning towards drowsiness or physical extenuation. Participants were instructed to associate an auditory stimulus (S) —high pitch sound or low pitch sound— with a response (R) button —left or right. In this auditory version of the PRL task, each S-R association leads to an auditory outcome (O)—correct (ding sound) or incorrect (white noise)— which participants use to assess their choice, and apply this knowledge to guide the next choices. Indeed, participants were explicitly told that there was a rule connecting each of the auditory sounds to a corresponding button (e.g., the low pitch sound could correspond to the left button, and the high pitch sound to the right button or vice-versa), which they had to figure out based upon instructive feedback they would receive after each R. Additionally, they were instructed on two key issues: 1) the S-R rule might switch after a certain amount of time —becoming the opposite of what it was previously— and that no specific indication whether such a switch had occurred would be provided; 2) although the majority of the time the feedback would be truthful, sometimes it could be false and in essence mislead to them. Therefore, the task entails the use of, at least, two rules to success, as participants have to press a button after each auditory stimulus (i.e., automatic rule) and to use an integrated history of S-R-O associations to determine the correct S-R association (i.e., high order rule). Once participants reach 90% accuracy or greater on the latest 10 trials, the implicit abstract S-R association is reversed, and participants have to infer the new association from the feedback received. The number of responses needed to attain a reversal (RAR) of the abstract association is used as the main index of performance. We hypothesized^9^ that reduced arousal states would lead to reductions in behavioural performance compared to baseline arousal state; while heightened arousal states would lead to improved performance relative to baseline, but only to an optimal point (i.e., moderate arousal) after which the performance will be deteriorated with further increases in arousal level (see figure 1A). These hypotheses were formulated in line with the famous psychology inverted u-shaped law originally attributed to Yerkes and Dodson (1908)^2^ relating arousal modulation performance in complex tasks, but later more formally defined by Broadhurst (1958)^47^ and Brown (1961).^48^

**Figure 1.**
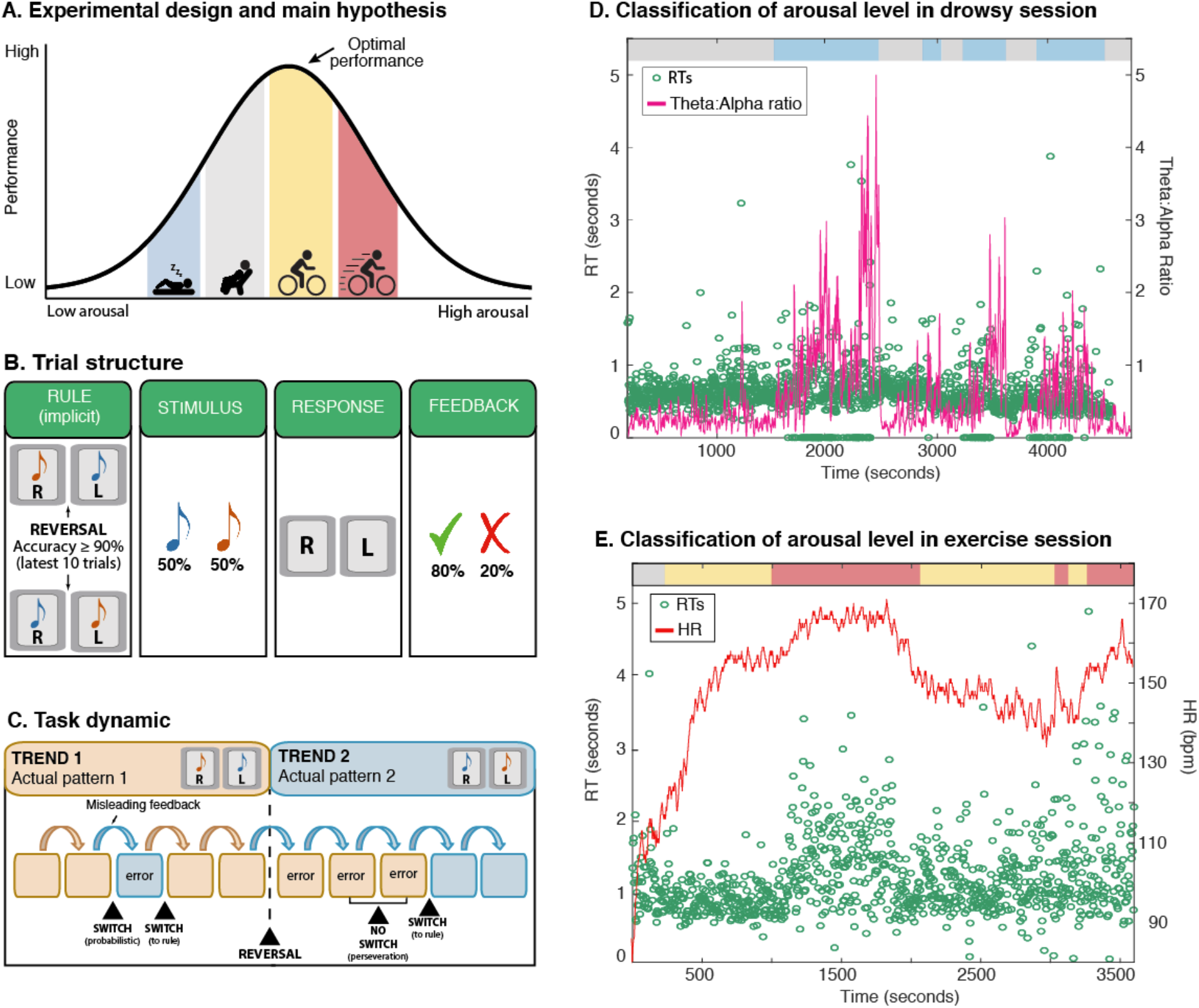
Experimental design and arousal level classification: A) Schematic representation of the experimental design and main hypotheses. Arousal level was endogenously manipulated by facilitating the natural transition of participants from awake to sleep, or instructing them to exercise during 60’ at the highest intensity and effort they could maintain without reaching premature extenuation. Notice that half of the participant transitioned towards drowsiness, while the other half transitioned towards physical exertion. A probabilistic reversal learning task was assessed continuously during the arousal modulation. Optimal performance of the task was expected at moderate arousal state (exercising at moderate intensity), while lower (drowsiness) and higher (exercising at high-intensity) arousal state were expected to result in task performance deterioration. B) In this auditory version of the probabilistic reversal learning paradigm, an auditory stimulus was presented on each trial, and participants had to associate the sound with a response button, left or right. After that, auditory feedback was provided according to the ongoing implicit rule. Notice that the feedback provided was not always truthful nor reliable, and attempted to mislead the participant 20% of the time. C) Task trials were grouped into sequences of trials following a particular rule (trend) where a particular sound was implicitly associated with a response button (e.g., high pitch sound with the left button, and low pitch sound with the right button). Participants were instructed to infer the rule from the provided feedback to assess their previous choice and apply the knowledge of their accuracy to guide the next choices, knowing that the rule might change after a certain time. Based on the feedback received, participants could make probabilistic or perseverative errors in the following trials. D) Automatic classification of arousal during a drowsy session (representative participant). The pink line depicts changes in the theta:alpha ratio (occipital electrodes cluster) during the pre-trial period (2 seconds before the auditory stimulus onset). The horizontal bars on top represent trials classified as baseline (grey) or low arousal (blue). The variability in the reaction times (green circles) closely follows the changes in theta:alpha ratio. Notice that circles on the horizontal axis (reaction time equal to zero) were non-responsive trials, usually during low arousal (drowsy) periods but also observed during exercise periods. E) Automatic classification of arousal during a physical exercise session (representative participant). The red line depicts changes in the heart rate during the pre-trial period (2 seconds before sound onset), and the horizontal bars on top represent trials classified as baseline (grey), moderate (yellow) or high arousal (red). Similar to the low arousal session, the reaction times (green circles) fluctuates with the changes in heart rate.

Note that, as a probabilistic task, the feedback provided is not always truthful nor reliable and misleads the participant 20% of the time (see figure 1B). Thus, the participant could correctly apply the S-R association and press the correct button in response to the auditory stimulus, and still receive negative feedback, thus indicating an incorrect choice. This scenario of conflicting evidence can lead participants to two different maladaptive response patterns (see figure 1C) while performing the task: 1) switching the pattern choice across trials with little (i.e., one negative feedback against the choice) or no evidence (i.e., no feedback against the choice) of an actual rule change (probabilistic switching); or 2) sticking with the previous choice despite having strong evidence (i.e., two or more negative feedbacks against the choice) of an actual rule change (perseveration). Relying on these response patterns lead to poor performance,^7^ as the optimal strategy in this task is to stick with the previous choice with zero or one negative feedback against the choice, and to switch the pattern choice if two or more consecutive negative feedbacks against the choice happen.

### Arousal modulates probabilistic information during a stream of conflicting evidence

First, we calculate the average RAR per participant in each arousal state (low, baseline sitting, baseline cycling, moderate, high). To account for the dependencies potentially generated by any procedural differences between Experiments, we fitted RAR using hierarchical linear mixed-effects modelling, with arousal as fixed effect, and participant nested into Experiment as random effects. The model showed a strong effect of arousal on RAR, F (3,113.02) = 11.59, *p* < 0.001, β = 0.61 (details on testing model assumptions can be found in the supplementary material), indicating that the processing of probabilistic information that allows the detection of changing patterns in a stream of conflicting evidence was modulated by the arousal level. Next, we checked for non-linearity in the relationship between arousal and RAR, to test the famous u-shaped curve. As expected, we found that the quadratic (*AIC* = 1243.6; *BIC* = 1262.3; *R*^2^ = 0.40) outperformed linear fitting (*AIC* = 1264.8; *BIC* = 1280.4; *R*^2^ = 0.23), confirming a possible curvilinear pattern (U shaped) of the effect of arousal on RAR (see figure 2), with a reliable increase in the number of responses required by the participants to complete a trend reversal (i.e., decrease of performance) as the level of arousal progress towards the extremes of the defined arousal range, confirming, for reversal learning, convergence with the Yerkes-Dodson law, later reformulated by Broadhurst in 1958.^47^

**Figure 2.**
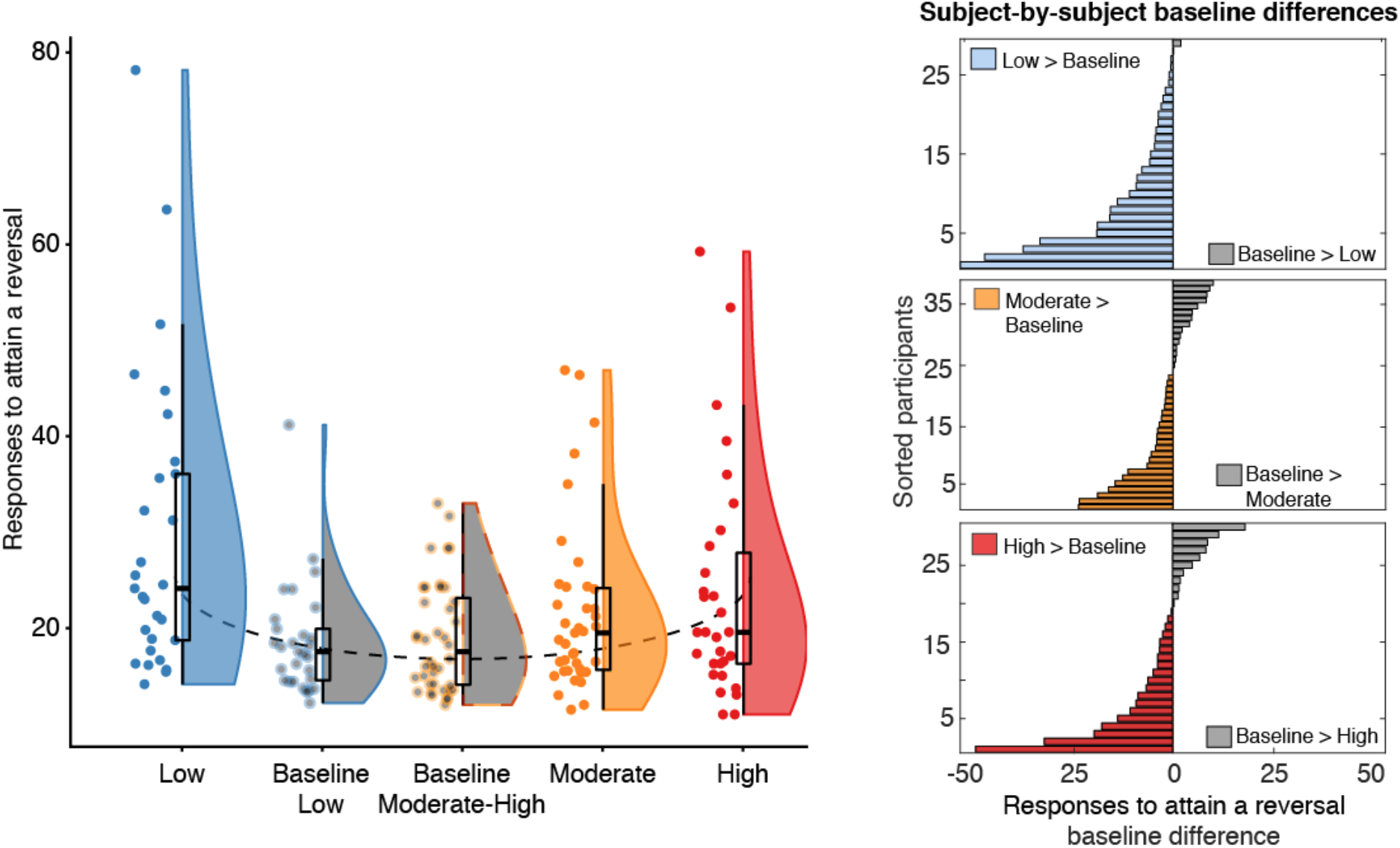
Number of responses needed to attain a trend reversal as a function of the arousal state. A) Violins and overlaid box plots of mean responses to reverse across arousal states. In box plots, middle black mark indicates the median, and bottom and top edges indicate 25th and 75th percentiles, respectively. The upper and lower whiskers indicate the maximum value of the variable located within a distance of 1.5 times the interquartile range above the 75th percentile and below the corresponding distance to the 25th percentile value. Surrounding the boxes (shaded area) is a rotated kernel density plot, which is comparable to a histogram with infinitely small bin sizes. Jittered dots represent the averaged response to reverse score for each participant in each arousal state. Linear mixed-effects model analysis revealed a reliable quadratic fitting between arousal and task performance, outlined by the dashed line. Low and high arousal states were associated with a worse task performance relative to their own baseline arousal states. Moderate arousal state was not associated with the expected optimal performance as no differences were found with the baseline arousal state. B) Baseline differences of each participant across altered arousal states are represented by the bars (grey bars indicate that these participants needed more trials to attain a trend reversal in the baseline compared with the altered arousal states; blue, yellow and red bars depict that these participants needed more trials to attain a trend reversal when arousal level was altered-increased or decreased-compared with baseline arousal state). Participants are sorted by performance difference between baseline and the arousal state. Upper and bottom panels show a consistent impairment of task performance across participants in low and high arousal states. Non-reliable differences were found between moderate and baseline arousal.

Splitting the comparisons to its specific baselines per arousal condition (i.e., sitting baseline compared to low arousal in the drowsiness condition; cycling baseline compared to moderate and high arousal in the exercise condition) yielded a reliable increase of RAR in low arousal, *t* (124.62) = 5.67, *p* < 0.001, β = 1.02, and high arousal state, *t* (117.93) = 2.57, *p* = 0.011, β = 0.45, compared with their corresponding baselines. Notably, baseline performance did not differ across arousal conditions (see supplementary figure 1). Contrary to what we expected, moderate arousal state was not associated with a decrease of RAR (the expected peak in performance), relative to baseline (*t* (114.85) = 1.61, *p* = 0.11, β = 0.25,). Moreover, we did not find evidence for a potential dual-task confounding effect in the heightened arousal conditions (see supplementary material). In sum, these findings provide evidence for an impairment in the processing of probabilistic information when the arousal level is altered, regardless of the side of the arousal spectrum.

### Different underlying mechanisms explain decreased performance in low and high arousal states

In the analysis above, performance under high and low arousal states was compared irrespective of the strategy participants may have used to solve the task. To test for the hypotheses of the differential mechanism driving changes in performance for each arousal side of the u-shaped curve, we calculated: a) probabilistic switching, as the proportion of trials when the participants change the pattern choice with little or no evidence (i.e., zero or one negative feedback against the choice); and b) perseveration, the likelihood of sticking with the previous choice despite strong evidence (i.e., receiving two or more negative feedbacks in a row) that the pattern has changed. Probabilistic switching and perseveration are proportion indices of strategic behaviour based on the probability of switching when negative feedback is provided. Thus, they range between 0 and 1, allowing comparison across arousal states while accounting for potential experimental differences (e.g., number of trials). We hypothesized that the impairment of performance in low arousal would be primarily attributed to an increase in probabilistic switching, relative to the baseline arousal state; and in contrast, the observed impairment of performance in high arousal state will be primarily due to an increase in perseverative behaviour. To test these hypotheses, we fitted probabilistic switching and perseveration (separately for low and high arousal states) using the hierarchical linear mixed-effects model structure defined previously. The analyses revealed that, while the probabilistic switching increased consistently across subjects during low arousal state compared with baseline arousal, F (1,56) = 14.78, *p* < 0.001, β = 1.01, *R*^2^ = 0.21, no reliable differences were observed in perseveration between these arousal states (F < 1). On the other hand, high arousal states led to a reliable increase in perseverative behaviour compared to the baseline state, F (1,67) = 9.12, *p* = 0.035, β = 0.34, *R*^2^ = 0.12, with no reliable differences observed in probabilistic switching (F < 1). These results suggest that altered arousal states lead to distinct maladaptive decision-making patterns that affect participants’ ability to generate stable evidence-based strategies, although evidence-driven responses were present (see figure 3A).

**Figure 3.**
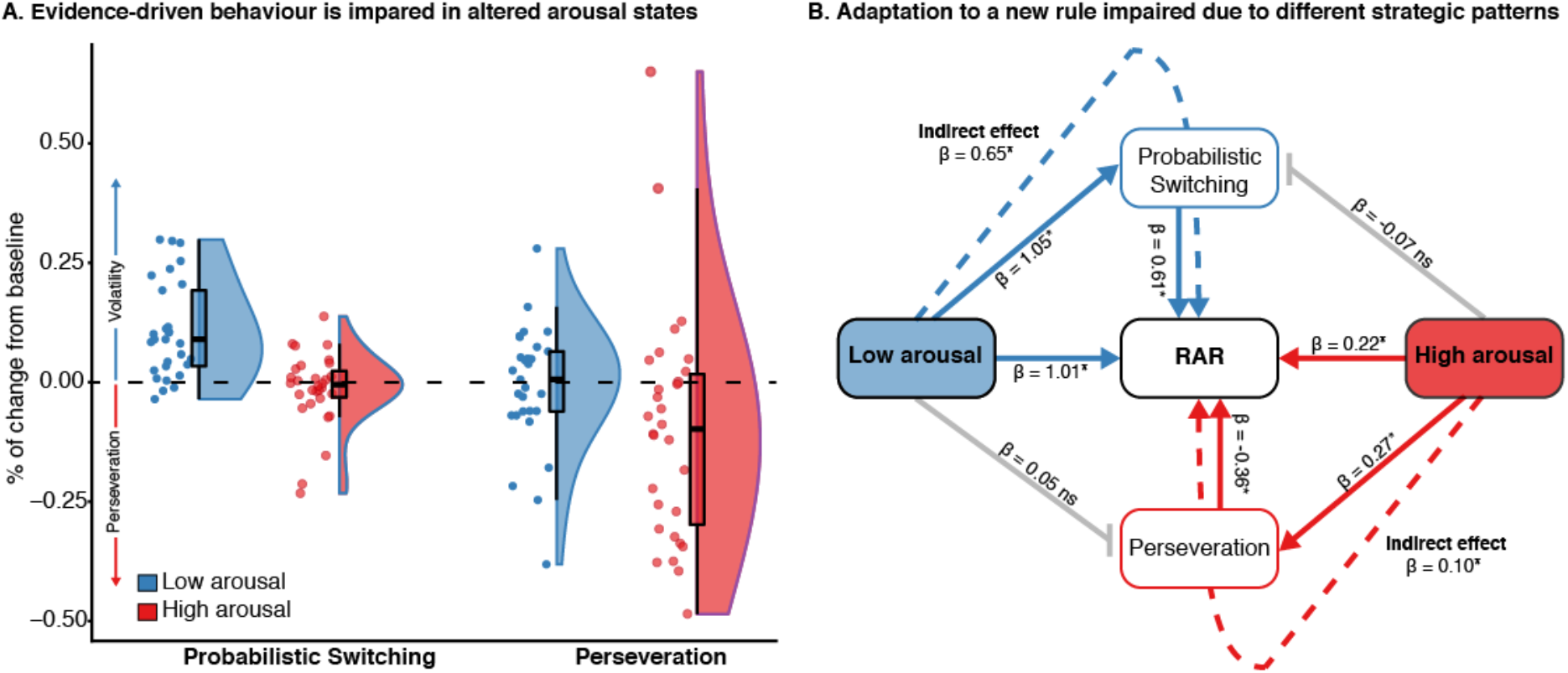
Maladaptive behavioural patterns across participants in low and high arousal states. A) Violins and overlaid box plots of the percentage of change from baseline to low (blue) and high (red) arousal states in probabilistic switching and perseveration. In box plots, middle black mark indicates the median, and bottom and top edges indicate 25th and 75th percentiles, respectively. The upper and lower whiskers indicate the maximum value of the variable located within a distance of 1.5 times the interquartile range above the 75th percentile and below the corresponding distance to the 25th percentile value. Surrounding the boxes (shaded area) is a rotated kernel density plot, which is comparable to a histogram with infinitely small bin sizes. Jittered dots represent the averaged response to reverse score for each participant in each arousal state. B) Mediation model diagram to illustrate that the general impairment in task performance found in low and high arousal states was mediated by different maladaptive behavioural patterns. Dashed lines (indirect effects) represent the effect of low (blue) and high (red) arousal on task performance (indexed by the averaged responses to attain a trend reversal) through probabilistic switching and perseveration, respectively. Solid lines depict direct effects between variables. Grey lines represent the absence of a direct effect of low arousal on perseveration and high arousal on probabilistic switching. Notice that a direct effect of an independent variable (arousal) onto the mediator (probabilistic switching, perseveration) is a prerequisite for mediation being possible. Standardized β regression coefficients are indicated in each effect (* depicts *p* < 0.05). Accordingly, the values of all effects are expressed as the number of standard deviations from the mean. For example, the direct effect of high arousal on RAR (β = 0.22) implies that a standard deviation change of 1 in the arousal variable would result in a standard deviation increase of 0.22 in RAR.

To further prove that the impairment in performance in low and high arousal states could be attributed to the different maladaptive behavioural patterns, we carried on a mediation analysis separately for each arousal state (low, high). We first confirmed that probabilistic switching and perseveration have an effect on the RAR, while controlling for the arousal state (see figure 3B). These results, together with the previous analyses where we found an effect of arousal state on probabilistic switching and perseveration, revealed a full mediation between these variables. As figure 3B illustrates, the regression coefficient between arousal and RAR, and the regression coefficient between probabilistic switching and RAR were statistically reliable, showing a full mediation of probabilistic switching on the effect of low arousal on RAR. The bootstrapped standardized indirect effect of low arousal on RAR, mediated by probabilistic switching, was 0.65 (*p* < 0.001), and the 95% confidence interval ranged from 0.29 to 1.07. A similar fully mediation effect was observed in high arousal state, showing that the effect of high arousal on behavioural performance was fully mediated via the perseverative behaviour. The bootstrapped standardized indirect effect was 0.10 (*p* = 0.014), and the 95% confidence interval ranged from 0.14 to 0.24. As predicted, participants showed an impairment of performance during low arousal state, relative to baseline arousal, which was primarily attributed to an increase of probabilistic switching (i.e., changing pattern choice with little or no evidence of an actual rule change). In contrast, while participants also showed an impairment of performance during high arousal state, relative to the baseline arousal, it was not attributed to an increase in probabilistic switching, but to an increase in perseverative behaviour (i.e., sticking with the previous choice despite consecutive negative feedbacks).

### Arousal disrupts the reversal strategy

To maximise performance in the task, a good strategy is to not fall for the false feedback and stand your ground until the next feedback, as well as switch to the second consecutive feedback. The fact that participants sometimes needed an unreasonable high number of responses to attain a reversal in low and high arousal states suggests the existence of sections of time on task in which they responded to the tones but could not apply the strategy rules (see fig 4A). These sections without clear strategic behaviour, that we call breakdowns, have been often neglected in previous studies using PRL tasks as failures of compliances or “bad participant”. The transient on/off nature of these breakdowns may provide valuable insight into the behavioural dynamics of participants in different states of arousal. We hypothesized that breakdowns sections would increase in low and high arousal states, relative to a baseline arousal state. First, we traced the sections of the task (more than 20 trials) in which participants did not attain a reversal. Second, we calculated the proportion of time these sections represented to the total time-on-task, and finally, we implemented a hierarchical linear mixed-effects model with the structure defined in previous analyses, separately for each arousal state (low, high), with the number of breakdowns as the index of performance. As hypothesized, low and high arousal states lead to longer breakdown sections compared with baseline arousal state (*t* (127.99) = 3.40, *p* < 0.001, β = 0.13; *t* (121.69) = −2.97, *p* = 0.003, β = 0.11). Subject-by-subject results (fig 4C) show a consistent increase of breakdowns across participants in low arousal state. Although high arousal states also showed a reliable increase of breakdowns as a group, this effect was less systemic, with half of the participants showing the opposite effect, no difference or no breakdowns.

**Figure 4.**
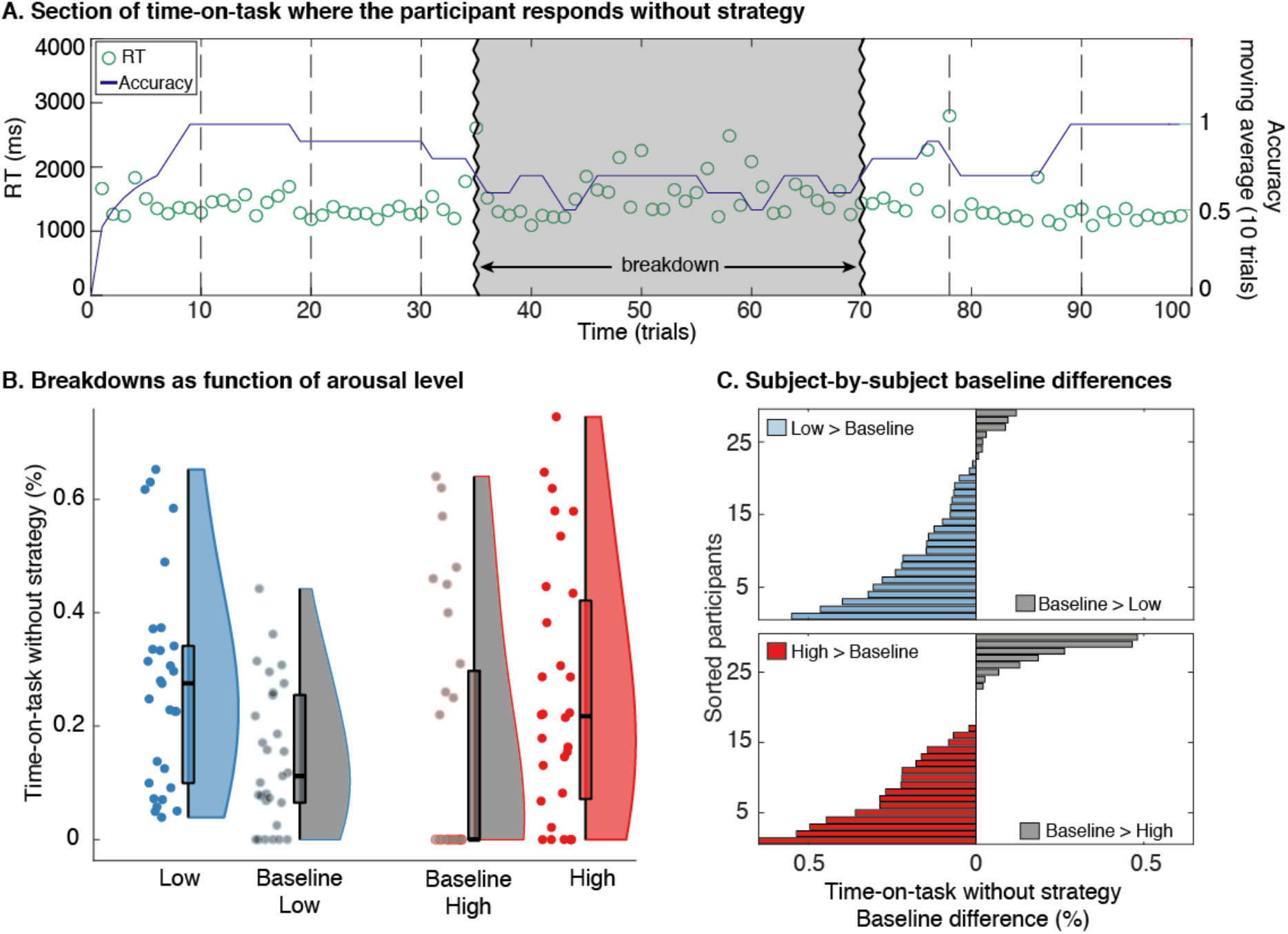
Behavioural strategy breaks as arousal changes. A) Automatic classification of a section of time where a representative participant responded without a clear behavioural strategy. The green circles show RTs and the blue line shows the ongoing accuracy of the task (10-points moving average). The grey shaded area flanked by the zigzagging vertical lines depicts the section of time classified as a breakdown. B) Violins and overlaid box plots of the averaged percentage of time-on-task without strategy across participants in low and high arousal states, compared with their respective baselines states. In box plots, the middle black mark indicates the median, and bottom and top edges indicate 25th and 75th percentiles, respectively. The upper and lower whiskers indicate the maximum value of the variable located within a distance of 1.5 times the interquartile range above the 75th percentile and below the corresponding distance to the 25th percentile value. Surrounding the boxes (shaded area) is a rotated kernel density plot, which is comparable to a histogram with infinitely small bin sizes. Jittered dots represent the averaged percentage of time-on-task without a strategy of each participant in each arousal state. Linear mixed-effects model analyses revealed that low and high arousal states lead to longer periods of breakdown relative to the baseline arousal state. Interestingly, violin plots show a considerable number of participants who had no breakdowns at baseline arousal states, something that completely disappears in low arousal state (all participants had breakdowns), and that is reduced in high arousal state. C) Baseline differences of each participant in low and high arousal states represented by horizontal bars (grey bars indicate that these participants spent more time performing the task without a particular strategy in the baseline arousal state compared with the altered arousal states; blue and red bars depict that these participants were applying behavioural strategies less time when arousal level was altered (increased or decreased) than in baseline arousal state. Participants are sorted by performance difference between baseline and the arousal state. Both panels show a consistent impairment of task performance across participants in low and high arousal states.

## DISCUSSION

In the present study, we facilitated natural transition of healthy participants towards the borders of non-pharmacological arousal states (drowsiness, physical exertion) to investigate the behavioural dynamics of cognitive flexibility. In line with our pre-registered hypotheses,^9^ the findings revealed a quadratic-like pattern (inverted U-shape) of the effect of arousal fluctuations on cognitive performance. As the level of arousal progressed towards the extremes of the defined arousal range reversal learning performance decreased, in agreement with the predictions of the Yerkes-Dodson law (1908).^2^ Although cognitive flexibility diminished in both under high and low arousal states, different maladaptive behavioural patterns drove this performance impairment. As predicted, the performance decline exhibited by our participants under drowsy states was primarily attributed to a more decision volatility (i.e., shifting pattern choice with little or no evidence of reinforcement contingencies change). In contrast, participants also showed a decline in performance during high arousal state but attributed to increased perseverative behaviour (i.e., sticking with a particular pattern choice despite having strong evidence that the contingencies have changed). Our findings also revealed that most participants undergo prolonged periods of time-on-task in which they seem unable to apply any specific higher order strategy. These breakdown periods, which can last for several minutes, are more frequent and sustained during high or low arousal. In short, our results provide solid evidence for distinct maladaptive decision-making patterns under altered arousal states, differentially affecting the participants’ ability to generate stable evidence-based strategies.

Arousal fluctuations thus seem to elicit a distinctive behavioural distortion of cognitive flexibility as further indicated by the microstructure of learning derived from trial-by-trial responses to negative feedback. Healthy participants under high arousal exhibited normal acquisition of S-R reinforcement contingencies but perseverative response patterns when contingencies were reversed. This failure to disengage from ongoing behaviour is a translational phenomenon strongly linked to impulsivity and compulsivity,^49^ and prevalent in numerous neuropsychiatric and medical conditions.^7,50,51^ For instance, patients with lesions that include ventral prefrontal cortex and orbitofrontal cortex,^52^ as well as chronic cocaine users^53^ and patients with schizophrenia,^54^ show normal acquisition of S-R contingencies but are severely impaired when those S-R reinforcement contingencies are abruptly reversed, exhibiting perseverative responding to the previously reinforced S-R contingency. Altogether, these findings suggest that high arousal undermines healthy individuals’ capacity to engage in complex cognitive strategies driving them to rely on habitual response patterns, which, paradoxically, might also enhance behavioural control in terms of response inhibition.^46^ Our findings not only further the understanding of the processes underlying automatized behaviour and habitual response tendencies, but high arousal may be used as a model to inform both impulsive and compulsive aspects of psychopathology.

In contrast, healthy participants under low arousal seemed unable to maintain the learned S-R reinforcement contingency and started to deviate from the evidence, revealing a volatile pattern of behaviour. Since a crucial aspect of the PRL experimental design was the existence of a 20% of misleading feedback, to maximise performance, individuals should not fall for the false feedback and —ideally— stand their ground until the next feedback. Further and as part of a successful strategy, they should switch if two or more consecutive feedbacks are given against the previously reinforced choice pattern. Consequently, adaptive behaviour during the task requires a balance between both types of behaviour (stability and flexibility). Those participants under low arousal fell repeatedly for the misleading feedback, switching prematurely after negative feedback. Furthermore, they showed increased decision volatility by spontaneously switching even without any negative feedback. This volatile pattern of cognitive flexibility has been linked to serotonin^55^ and dopamine systems,^56^ and is observed in patients with major depression,^57,58,59^ often linked to either an oversensitivity to punishment or an impaired control over negative feedback.^60,61^ It is reasonable to speculate that low arousal levels render individuals more sensitive in updating S-R reinforcement contingencies, rather than increase sensitivity to punishment as in major depression. Moreover, low arousal may increase volatility by decreasing attentional resources, leading to spontaneous explorations, higher RT variability and periodic omissions (see supplementary figure 2).

The fragmentation of cognitive control due to changes in arousal has been primarily shown in sleep deprivation^62,63,64,65,66^ and not in spontaneous fluctuations of alertness as we show in this study. The increased volatility in the PRL with low arousal suggests a decrease in cognitive control that is different from an increase in perseverative behaviour seen in high arousal. Indeed, we have previously shown that decreased levels of arousal can fragment or reconfigure specific aspects of cognition while preserving crucial executive control processes such as the capacity to detect and react to incongruity,^18^ the efficiency in perceptual decision making,^31^ and the precision of conscious access.^17^ Here, we add further evidence showing that individuals under reduced arousal state, although struggling to maintain stable evidence-based decision-making patterns, are able to learn new S-R reinforcement contingencies, demonstrating flexibility of the human brain to adapt to increasing levels of endogenous (arousal) noise. The evidence of cognitive and —indirectly— neural reconfiguration of cognitive control networks suggests compensatory mechanisms elicited by the change in arousal.

Upon further examining the microstructure of learning derived from trial-by-trial performance of the PRL task, we uncovered the existence of prolonged periods of time-on-task in which participants did not seem to apply any particular high-order behavioural strategy. Although these breakdown periods emerged regardless of the arousal level, they were prevalent under low and high arousal states, lasting from few to several minutes. Remarkably, the transient on/off nature of these breakdowns suggests that extreme arousal levels alternate between different metastable cognitive states. The first state can be defined by a relatively successful application of the reinforcement information where participants can navigate the uncertainty of the PRL, while in the other metastable state they seem to only apply the simple auditory-motor S-R rule to respond to the auditory tones but are unable to use choice history to develop a successful strategy.

In the context of this study, arousal as a biological construct defined by the homeostatic regulatory capacity of the system and its responsiveness,^67^ helps to link drowsiness and increased alertness during physical exercise in a common framework where the predictions of the Yerkes-Dodson inverted U-shaped law can be experimentally tested. Despite the obvious difference at the biological, neural and psychological level between both sides of the curve, the common decrease in performance highlights the commonalities between the extremes in human performance, adding to the fact that both —sleep and physical exertion— emerge as natural transitions from a similar state (resting) traversing different stages, and exhibit nonlinear dynamics and hysteresis processes in their transitions.^12^ Thus, drowsiness and physical exertion provide complementary perspectives on cognitive dynamics when the arousal level is altered. The present findings point out their differences in the cognitive fragmentation leading to a general decline in task performance.

Transitions towards drowsiness or physical exertion entail changes in levels of arousal, which are in turn associated with a wide range of alterations (e.g., neural, motor, endocrinal, phenomenological, etc.) that might cause the cognitive fragmentation described in the present study. For instance, during a single bout of aerobic exercise, as intensity increases from low to high, there is a release of epinephrine and, to lesser extent norepinephrine, into the blood from the adrenal medulla.^15^ This exercise-induced increase in brain concentrations of catecholamines has been proposed as a physiological mechanism underlying cognitive performance during and after physical exercise.^15^ Similarly, when falling asleep, we experience a cascade of changes in almost every system of the organism, including the somatic and autonomic nervous systems,^12^ which might be playing a crucial role in cognitive processing. The extent to which each of the changes that occur during these transitions (drowsiness and physical exertion) are responsible for the cognitive adaptations we report here is something that future studies might reveal, for example, combining measurements of the autonomic nervous system and brain functioning, which would make it possible to gain more insight into the underlying physiological mechanisms involved in arousal-related changes in cognition. These inferences of this study are hence mediated by physiological processes that might partially explain the cognitive modulations in an independent manner if dissociated from arousal changes.

Though the Yerkes-Dodson law was not initially formulated to be a general rule to apply to all psychology subfields (learning, motivation, emotion, etc.), through the years, and with the pressure to find common mechanisms in psychology, the findings initially defined for learning were further extended and reinterpreted as a law about the relationship between arousal and other physiological constructs to perceptual and cognitive performance.^68^ Despite this overgeneralization from its genuine formulation and its reductionist nature, our findings rely on such inverted U-shaped law as a basic useful theoretical framework, providing an attractive theoretical model to characterize the neural, cognitive and behavioural dynamics involved in the impact of arousal fluctuations in a wide range of physiological states and neuropsychiatric conditions.

Our findings bring some generalizations about the need to extend the traditional framework of understanding the interplay between cognitive dynamics and arousal through the prism of the homeostatic steady-state dynamics using pharmacological interventions^34^ or transient alterations of emotional state.^69^ In addition to this classical approach, we believe that drowsiness and physical exertion provide fruitful —naturally occurring— alterations of the arousal level with a preserved capacity to behaviourally respond, which can be utilized to study the modulation of neural function and cognitive processing. In the traditional steady-state approach, such natural fluctuations of the arousal level may be undetected,^70^ hindering or distorting cognitive and neural markers of crucial aspects of information processing.^17^ Pharmacological and lesion perturbations of the brain are regarded as causal in cognitive neuroscience and regarded as stronger in their explanatory power than conditions relying on stimuli or psychological modulations. Arousal is an internally modulated change that can be used to study cognition and may be regarded in the strong causality range due to its partial independence from psychological processes.^18^ The cases of drowsiness and physical exertion as causal models to study the neural mechanism of cognitive flexibility may prove to be very useful in the exploration of how cognition is fragmented or remain resilient under (reversible) perturbations of arousal^17,33,71^ Our findings highlight that further research should focus on the rapidly changing dynamics of brain function and cognitive processing that appear to capture key dynamics relevant to our behavioural and perhaps even phenomenological experience, as we drift into strained physiological states.

## MATERIALS AND METHODS

### Participants

A total sample of 100 participants of an age range between 18 and 40 years old was included in the present study. All participants reported normal binaural hearing, no visual impairment and no history of cardiovascular, neurological or psychiatric disease. They were asked to get a normal night rest on the day previous to testing, and not to consume stimulants like coffee or tea on the day of the experiment.

The first experiment (herein Experiment#1) consisted of 35 participants (15 female; age range 18-40). In addition to the general aforementioned inclusion criteria, only easy sleepers, as assessed by the Epworth Sleepiness Scale (ESS),^72^ were selected to increase the probability that participants fell asleep. Recruited participants were considered healthy with relatively high ESS scores but not corresponding to a condition of pathological sleep such as hypersomnia (i.e., scores 7–14). They were recruited via the Cambridge Psychology SONA system. Note that the target sample size was 50 participants transitioning towards drowsiness. However, after collecting the first 35 we decided to make slight modifications to the experimental protocol by increasing the time of the drowsy blocks to obtain a higher proportion of trials in low arousal. For this reason, we decided to collect a second sample (Experiment#2) which consisted of 15 participants (11 female; age range 18-40), where we included these key modifications to the experimental protocol (see Procedure section for more details). Inclusion criteria and recruitment processes were similar to Experiment#1.

The third experiment (herein Experiment#3) consisted of 50 participants (6 female; age range 19-39). Additionally to the common inclusion criteria, only individuals who reported at least 8 hours of cycling or triathlon per week were selected. Well-trained cyclists were selected because they are used to maintaining the pedalling cadence at high intensity during long periods of time. Furthermore, they are able to keep a fixed posture over time, which notably reduces movement artefacts. They were recruited from the University of Granada (Spain) through announcements on billboards and previous databases.

All participants from the three experiments gave written informed consent to participate in the study and received a remuneration of 10€ per hour (i.e., approximately 30€ per participant). The Cambridge Psychology Ethics Committee and the University of Granada Ethics Committee approved the study (CPREC 2014.25; 287/CEIH/2017).

### Experimental task

A modified version of the probabilistic reversal learning paradigm was used in all three experiments, which was characterized by employing auditory stimuli and an abstract rule (see figure 1B-C). In this task, participants learnt to choose one of two randomly presented tones by receiving instructive auditory feedback tones after each response, indicating either a correct or incorrect choice. When participants reached a 90% accuracy in the last 10 trials, reinforcement/punishment contingencies were reversed so that the previously reinforced tone was punished and vice versa. Within each reversal trend, a 20% probabilistic error trial was included in which “wrong” feedback was given for correct choices, even though the reinforcement contingencies had not changed. Participants were instructed to infer the rule from the feedback received, knowing that sometimes it might be misleading and that the rule might change after a certain time (see supplementary material for more details on the task instructions). The stimuli were binaurally presented at a random time interval (between 1000 and 1500 ms) during 500 ms. They had to respond to both targets by pressing a button with their right or left hand.

### Procedure

In Experiment#1, participants were fitted with an EGI electrolyte 129-channel cap (Electrical Geodesics, Inc. systems) after receiving the task instructions and subsequently signing the informed consent. The whole session was completed in a comfortable adjustable chair with closed eyes. Task instructions were to respond as fast and accurately as possible, reducing body movements as possible and keeping the eyes closed. In the beginning, the back of the chair was set up straight and the lights in the room were on. Participants were asked to remain awake with their eyes closed whilst performing the first block (awake block) of the task which consisted of 480 trials, lasting 30 min approximately. Then, the chair was reclined to a comfortable position, the lights were turned off and participants were offered a pillow and a blanket. They were explicitly told that they were allowed to fall asleep during this part of the task and that the experimenter would wake them up by making a sound (i.e. knocking on the wall) if they missed 5 consecutive trials. This block (drowsy block) also consisted of 480 trials. Then, the sequence of two blocks (awake-drowsy) was repeated. In total, participants completed 1920 trials divided into 4 blocks of 480 trials each one. The whole session lasted for 3 hours approximately.

In Experiment#2, the procedure was similar to the Experiment#1 except for the time to fall asleep that was increased to get a higher amount of low-arousal (i.e., drowsy) trials. Participants completed a total of 2120 trials, divided into 4 blocks. The order of the blocks was the same for all participants and followed the same sequence as in Experiment#1: awake-drowsy-awake-drowsy. Awake blocks had 100 trials each one, while drowsy blocks consisted of 960 trials each one. The session lasted for 3 hours approximately.

In Experiment#3, upon arrival to the laboratory, participants were seated in front of a computer in a dimly illuminated, sound-attenuated room with a Faraday cage. They received verbal and written instruction about the experiment and were prepared for electrophysiological measurement. They were fitted with a 64-channel high-density actiCHamp EEG system (Brain Products GmbH, Munich, Germany) and a Polar RS800CX heart rate (HR) monitor (Polar Electro Öy, Kempele, Finland). Notice that EEG data was acquired but was not used to test the hypotheses of this study, and will be reported elsewhere. The whole session consisted of 4 different blocks. The first one was an adaptation (non-exercise) block in which participants performed 100 trials while resting in a comfortable chair. Then, they got on a cycle-ergometer and completed 100 trials while warming-up at light intensity. Subsequently, they completed a self-paced 60’ time-trial (i.e., high-intensity exercise) while performing the task, resulting in 850 trials approximately (the number of trials slightly varied as a function of the reaction time of participants). In line with previous experiments from our laboratory,^73,74,75^ in the self-paced time-trial participants were instructed to achieve the highest average power (watts) during the 60’ time-trial exercise, and were allowed to modify the power load during the exercise. They were encouraged to self-regulate effort in order to optimize physical performance without reaching premature exhaustion. That self-regulation yielded fluctuations of effort during the 60’ exercise period, which allowed us to study the effect of arousal on the management of probabilistic information. Once the 60’ time-trial block was finished, participants completed the last block while cooling down at light intensity, which was also composed of 100 trials. All participants completed the blocks in the same order, lasting around 3 hours.

### Arousal classification

To capture the arousal fluctuations during the transitions towards drowsiness or physical exertion at the single-trial level, we implemented two different analytical approaches which were pre-registered after data collection.^9^

In Experiment#1 and Experiment#2, the arousal level was endogenously manipulated by facilitating the natural transition from awake to sleep. This transition reduces arousal and yields a considerable proportion of drowsy yet responsive trials as seen in previous experiments from our laboratory.^17,30,71^ This way, we were able to study the effect of arousal (i.e. baseline arousal [awake] trials vs. low-arousal [drowsy] trials) on the management of probabilistic information. Given that awake-sleep transition is characterized by a decreasing alpha range activity, together with an increasing theta range activity (Hori et al., 1994), progression of drowsiness was quantified by the spectral power of respective EEG frequency bands ^i^. We computed the spectral power of EEG frequency oscillations for each trial from −2000 ms to 0 ms in respect to the onset of a target tone using continuous wavelet transform, set from 3 cycles at 3 Hz to 8 cycles at 40 Hz. Theta (4-6 Hz) and alpha (10-12 Hz) power were then averaged individually for each trial across central (E36, E104) and occipital (E75, E70, E83) electrodes for theta and alpha rhythms respectively. Finally, theta/alpha ratio was computed and smoothed with a 4-point moving average resulting in a single “sleepiness” value per trial. Visual inspection of theta/alpha ratio and RT dynamics of each participant confirmed the presence of clear sleepiness-related fluctuations during the experimental session, especially during drowsy blocks. Those participants who did not show clear fluctuations of the theta:alpha ratio were removed from final analyses (5 subjects). Then, each trial for each participant was initially categorized as drowsy (top 33% of lower theta-upper alpha ratio scores) or alert (lowest 33%). Further, following the sleep hysteresis physiology criteria^77^ isolated awake trials within prolonged periods of drowsy (≥10 trials) were considered as drowsy to account for the gradual homeostatic change during the sleep transition. In addition, the first 100 trials of each block (awake and drowsy) were considered as awake trials.

In Experiment#3, the arousal level was endogenously manipulated by facilitating the natural transition from a resting state to high-intensity physical exercise. This transition increases the arousal level progressively, with continuous fluctuations that affect cognitive performance as seen in previous studies from our laboratory.^40,75,78,79^ We captured these arousal fluctuations at a single trial level (moderate arousal trials, high arousal trials) by using the HR response. To address the intersubject variability, HR data were transformed into differential scores relative to the HRmax estimated using the equation of Tanaka et al., (2001)^80^, a reliable and well-established method to calculate HRmax in healthy individuals. Then, moderate and high arousal trials were characterized based on percentage relative to HRmax. HR between 60% and 80% of HRmax were considered as moderate arousal, while HR higher than 80% HRmax were considered as high arousal. Due to technical issues with HR monitoring, 4 subjects were removed for further analyses.

### Behavioural data analysis

In probabilistic reversal learning paradigms, participants are instructed to infer an abstract rule form the feedback they receive, knowing that sometimes it might be misleading and that the rule might change. Since a reversal is triggered when a high-level accuracy is reached, the number of responses needed to attain a reversal is considered one of the main indices of performance. To delineate the microstructure of learning derived from trial-by-trial responses we considered the likelihood of switching the pattern choice across trials as a function of the amount of consecutive negative feedback received. The likelihood of switching was considered the main index of strategic behaviour, and was divided into 2 different strategies: *i*) Probabilistic switching: the proportion of trials when the participants change the pattern choice with little (one negative feedback against the choice) or no evidence (no feedback against the choice) of an actual rule change; *ii*) Perseveration: likelihood that participants stay with the seemingly incorrect choice even after receiving two or more negative feedbacks in a row).

The number of breakdown sections was also used as an index of performance. We defined a breakdown as a section of time in which participants ‘lose’ the task, and do not follow any strategy, being unable to reach a change of trend during more than 20 consecutive trials. RT, accuracy, and omissions were also checked as secondary indices of behavioural performance.

Participants with overall accuracy under 60% or less than 3 reversals attained during the baseline period were excluded (i.e., 4 subjects from Experiment 1; 2 subjects from Experiment 2; 6 subjects from Experiment 3).

### Statistics

#### Single-subject analysis

In order to test the hypotheses, we took a set of strategies. We first captured the direction of effects for each of the key performance variables (i.e., RAR, RT, accuracy, omissions, and switching likelihood), and contrasted them for each participant, obtaining an indication of the direction and strength of the effects per participant. Descriptive and distribution measures, as well as single-subject statistics, were used as guidance of the variability of effect size in single variables, and for guiding the previously defined exploratory hypotheses. Per participant, effect sizes were calculated and depicted for each of the key performance variables to check the effect size of individual differences across arousal states.^ii^

#### Group analysis

To investigate the management of probabilistic information as a function of arousal, we conducted mixed-effects analyses including data from the three experiments collapsed into a single dataset with RAR as the main index of performance. In face of the diversity of samples’ characteristics and experiment features, we fit RAR using hierarchical linear mixed-effects modelling, as implemented in the lme4 R package.^81^ We treated RAR as obeying to a hierarchical data structure with arousal as fixed effect, and participant (level 2) nested into experiment (level 1) as random effects. This random part was common to all models. We tested the specific hypothesis by using the same approach based on multilevel linear mixed-effects modelling. Different variables (i.e., probabilistic switching, perseveration, breakdowns, RT variability and omissions) were analysed in a multilevel data structure, with the fixed (arousal) and random effects (experiment/participant) adjusted to the specific hypothesis tested.

Models were compared using the Akaike Information Criterion (AIC), and a likelihood ratio test. Notice that AIC does not assume that the true model is among the set of candidates (and is just intended to select the one that is closest to the true one). In our case, fitting decisions were not about the truthiness of models, but to include or not a given factor. For model comparisons performed to identify the best-fitting model, a relatively lenient 0.010 p-value criterion was adopted.

Causal mediation analyses were conducted to estimate the proportional direct and indirect effects of arousal on task performance through probabilistic switching and perseveration strategies (mediators) using the “mediation” package in R^iii^.^82^ This method allowed us to assess a confidence interval of the mediation effect itself using rigorous sampling techniques with fewer assumptions of the data. The average causal mediation effect was determined using a nonparametric bootstrapping method (bias-corrected and accelerated; 1000 iterations) and reported as standardized β regression coefficients for direct comparison with each other. Confidence intervals were obtained using a quasi-Bayesian approximation.

## Pre-registration

The hypotheses and analyses plan were pre-registered in the OSF repository after data collection (https://osf.io/tzw6d).

## Data and code

Data and codes used for the analyses presented here are available at the OSF repository (https://osf.io/xk379/).

## Funding

This research was supported by a University of Granada Postdoctoral Fellowship (2019/P7/100) and the Junta de Andalucía (DOC_00225) to L.F.C, a research grant from the Ministerio de Economía y Competitividad (PSI2016-75956-P and PID2019-105635GB-I00) to D.S., and Wellcome Trust Biomedical Research Fellowship (WT093811MA) awarded to T.A.B.

## SUPPLEMENTARY MATERIAL

### Dual-tasking effect of physical exercise on baseline performance

Is the detrimental effect of heightened arousal on behavioural performance truly due to the increased arousal level, or does it simply reflect a dual-task confounding effect of the physical and the cognitive task occurring simultaneously? Although this question is partially tackled in main analyses as the baseline arousal state of the heightened arousal states was also a dual-task condition (i.e., warm-up), we specifically explored whether a dual-tasking arousal baseline might be associated with poorer performance (i.e., higher RAR and RT variability), relative to a non-exercise adaptation period that participants performed just before the warm-up. Contrary to what we expected, the mixed-effects model yielded no reliable performance differences between the adaptation period and the warm-up (RAR: *t* (39) = 1.41, *p* = 0.167, β = 0.18; RT variability: *t* (39) = 1.50, *p* = 0.14, β = 0.19). To further confirm that baseline performance was equal or similar for all experiments, we analysed the behavioural performance during baselines of Experiments 1 and 2 (i.e., wakefulness periods), as well as during baseline of Experiment 3 (i.e., warm-up period). Neither the number of responses needed to attain a reversal (RAR) nor accuracy showed reliable differences in baseline performance between experiments (F < 1). Subject-by-subject results show a similar distribution of performance across subjects in each Experiment (see supplementary figure 1).

**Supplementary figure 1:**
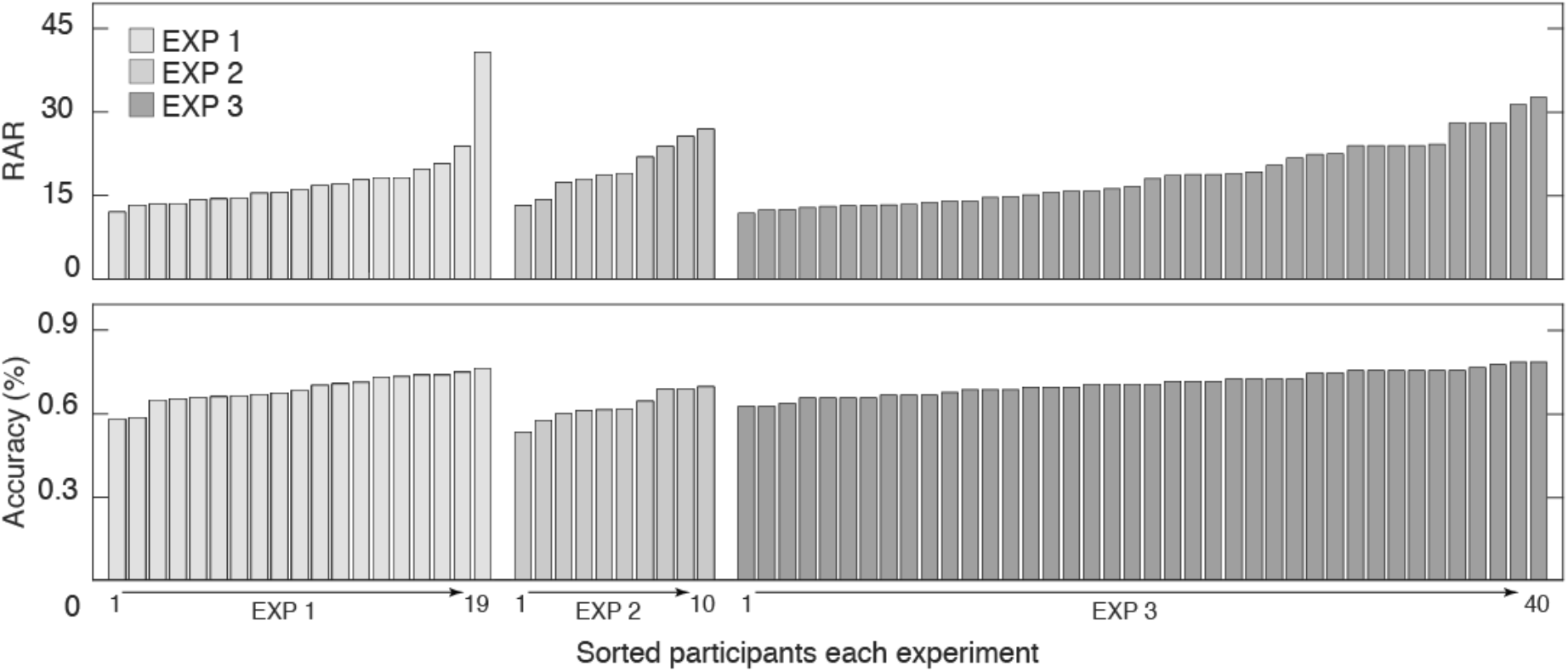
Subject-by-subject baseline performance. Individual behavioural measures during baseline across databases. Grey bars represent individual participants within each experiment. All subjects are arranged by performance, from best to worst in RAR, and from worst to best in accuracy. The analysis revealed no reliable differences in behavioural performance during baseline periods across experiments.

### Low arousal deceleration in behavioural dynamics

The transition from wakefulness to sleep involves a progressive, and sometimes nonlinear loss of responsiveness to external stimuli and a progressive increase of RT variability.^10,18,30^ To further characterise the behavioural pattern of this transition, and compared to previous falling asleep tasks, we investigated the responsiveness and RT dynamics of the participants in the low arousal condition. We fitted a mixed-effects model separately for RT variability and omissions as dependent variables. As predicted from other cognitive tasks, ^17,18,30,33^ low arousal led to higher RT variability (*t* (27.99) = 4.59, *p* < 0.001, β = 0.54), which was accompanied by a drastic increase in omitted response to stimuli (*t* (27.99) = 5.11, *p* < 0.001, β = 0.67), compared with the baseline arousal state (see supplementary figure 2). These findings confirm the convergence to other tasks of our arousal manipulation in probabilistic reversal learning in its basic effects.

**Supplementary figure 2.**
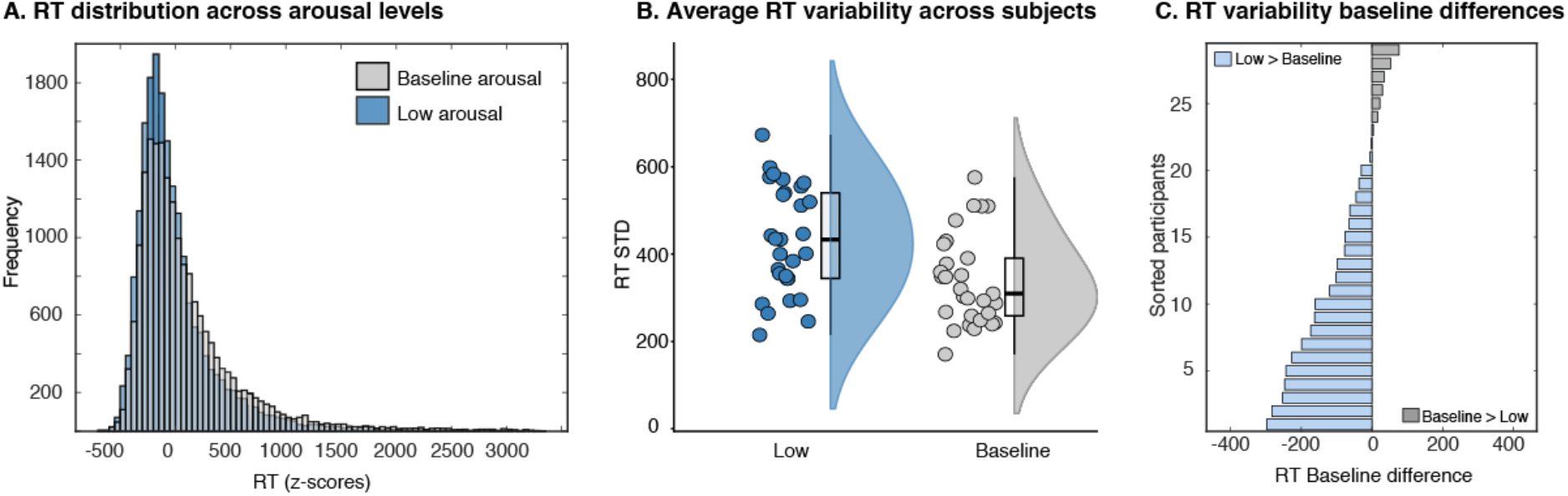
RT dynamic in low arousal. A) RT distribution during low (blue bars) and baseline (grey bars) arousal states. B) Violins and overlaid box plots of the averaged reaction time variability across participants in low and baseline arousal states. C) Subject-by-subject baseline differences in RT variability in low arousal. Grey bars represent participants with a higher RT variability in the baseline compared with low arousal state. Blue bars depict participants with a higher RT variability when arousal level was reduced compared with baseline arousal state. Participants are sorted by the RT variability difference between baseline and the arousal state.

### Testing the assumption of the final mixed-model

We tested the main assumptions of mixed-models on the hierarchical linear mixed-effects model (m1) we used to assess the effect of arousal of RAR with participant nested into Experiment as random effects. We first tested the linearity of the data by plotting the model residuals (i.e., the difference between the observed value and the model-estimated value) against the predictor (see supplementary figure 3A-B). As we predicted, the relationship between arousal and RAR is not well described by a straight line (Shapiro-Wilk normality test = 0.85963, *p* < 0.001) but by a quadratic function as shown in the Results section. Then, we checked that the covariance of the residuals was equal across experiments and participants. We run a variation of Levene’s test by calculating the absolute value of the residuals from the model, and squaring them for a more robust analysis with respect to issues of normality (Glaser 2006). Neither the ANOVA of the between experiments residuals (F < 1) nor the ANOVA of the between subject residuals (F = 1.24, p = 0.163) yielded significant differences. Therefore, our model met the assumption of homoscedasticity. Finally, we estimate whether the residuals of the analysis were normally distributed (see figure 3C). The QQ plot provides an estimation of where the standardized residuals lie with respect to normal quantiles, showing a light deviation from the provided line that suggest that the residuals themselves were not normally distributed. The violation of this normality assumption however has been proven that do not noticeably impact results where the number of observations per variable is higher than 10 (Schmidt & Finan, 2018).

**Supplementary figure 3.**
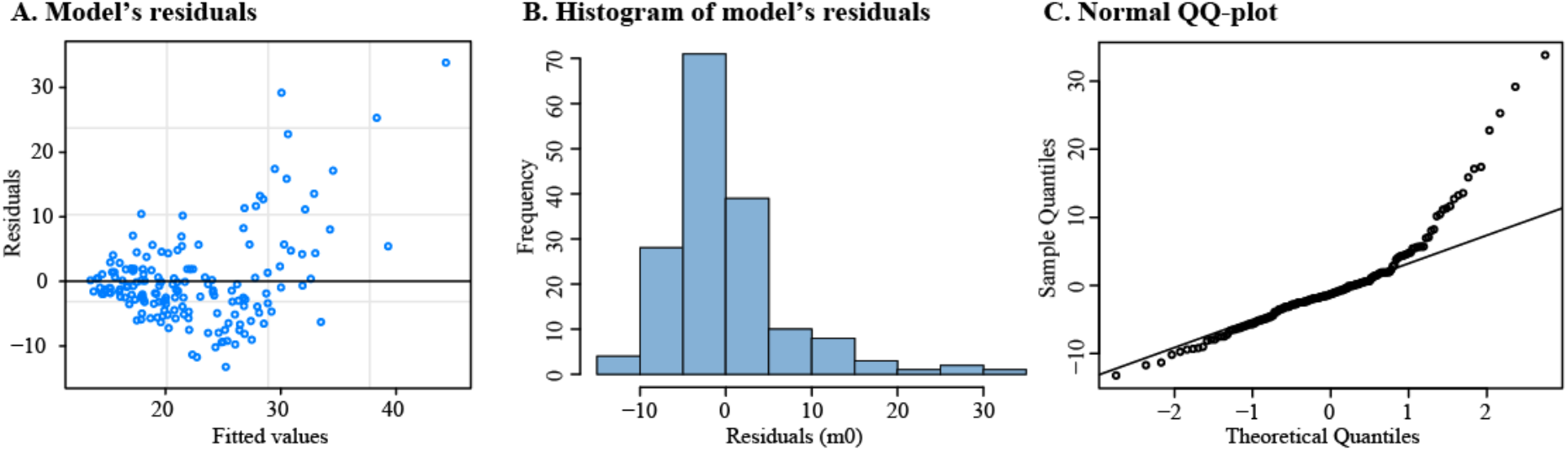
Model validation graphs. A) Fitted values versus model’s residuals (homoscedasticity). B) Histogram of model’s residuals (normality). C) Estimation of the linearity of the residuals (linearity).

### Probabilistic reversal learning task instructions for participants

This experiment will consist of four separate blocks. First there will be 2 short ‘alert’ blocks (5-10 minutes in length) during which you will remain fully attentive and try your best at completing the task. Then afterwards there will be 2 longer ‘drowsy’ blocks (45-60 minutes in length) during which you still need to complete the task, however the lights will be turned off, your seat reclined, and you can relax and embrace whatever drowsiness/boredom comes along.

The task itself involves listening to sounds through a set of headphones and then responding to those sounds with the button box provided. There are two separate sounds that you can hear, either a low pitch sound or a high pitch sound, and there is a rule that connects each of these sounds to a corresponding button on the button box. For example, the low pitch sound could correspond to the left button, and the high pitch sound to the right button, or vice versa. You will not know what the rules are when you first begin the task, but rather you must figure them out based upon instructive feedback that you will receive after each response, indicating either a correct or incorrect choice. If after hearing the stimulus sound you correctly press the corresponding button, you will hear a nice ‘*ding’* sound as feedback indicating you made the correct choice. However, if your response on the button box is not in line with the current rule, you will hear a ‘*static’* noise as feedback, indicating an incorrect response.

At this point ask the participant if they fully understand how their job works for the task. Feel free to elaborate and go over it again and again if necessary. There is little risk of influencing the data based upon variations in instruction for this core aspect of the experiment, and it is imperative that they fully understand this basic part otherwise their data WILL actually be corrupted. Additionally, if they do not fully understand up until this point, they will certainly become even more confused with the remaining information.

Now because we like to make things complicated, there are two important caveats to remember in terms of this experiment. The first is that the rule that connects each sound (high or low) to a specific button (left or right) will not always remain the same. After a certain amount of time the rule may switch and therefore become the opposite of what it was previously. So for example, if previously the rule was the high sound corresponds to the left button and the low sound to the right button, the new rule would be high equals right and low equals left. This switching of the rules can occur more than one time throughout each block, and you will receive no specific indication if a switch has occurred. It is up to you to figure out if and when a switch happened based upon your choices and the feedback you receive.

The second caveat is that although **the majority of the time** the feedback you receive in response to your button choice will be truthful, sometimes it will be false and in essence lie to you/try to trick you. So for example, if the current underlying rule states that the high pitch sound corresponds to the left button, and you make the correct choice (press the left button after hearing the high pitch sound), the feedback **could** be the *‘incorrect static’* sound. Again, most the time the feedback will be truthful and not trying to deceive you, but it is important to keep in mind that it can happen occasionally.

If the participant has any questions after explaining these caveats, be mindful of how you answer. Feel free to go over them again, but try to basically use the same wording as the first time you explained. This is because small differences in word choice have the potential to greatly influence how the participant approaches the task (how often they feel the rules switch, how much they trust the feedback, etc.).

Now because this can seem quite complicated at first, you can have a quick practice session to get used to the experiment. This is often helpful to participants and will hopefully make you more comfortable in your understanding of the task. If you still have any questions afterward we can briefly go over it again.

If after the practice session the participant still seems to not understand the task, you can go over it again but still be mindful of your word choice. There is a difficult line to straddle between making sure the participant fully understands the task, and making sure we are consistent in our explanation and therefore the participants’ approach. Straying too far in either direction can greatly influence the data.

After the practice session ends it is time to begin the first ‘alert’ block of the experiment.

Now we are going to begin the actual experiment. First up we will have the two short *‘alert’* blocks which will be around 5-10 minutes each. When performing the task during these two blocks please remain fully alert and focus on the task as much as possible. Your eyes must remain closed during the duration of these blocks.

After the completion of the two ‘alert’ blocks, get the participant comfortable for the ‘drowsy’ portion. It is very important to make the participant as comfortable as possible. This can include reclining their chair, supplying pillows or blankets, participants removing their shoes, etc. It is VERY important they are actually comfortable (and not just saying “yeah sure whatever”).

Now we are going to begin the ‘drowsy’ portion of the experiment. There will be two separate blocks, around 45-60 minutes each. You are encouraged to become as comfortable and relaxed as possible, and to embrace any feelings of drowsiness that come over you. However, you do of course still need to try to complete the task so you cannot sleep throughout the entire experiment. If I see that you have actually fallen asleep, I will lightly knock on the door to wake you back up, no worries! I will turn the lights off for these two blocks and once again please keep your eyes closed for the duration of the blocks.

## Footnotes

i Deviation from pre-registration. Originally, we aimed to use the automated offline method developed by Jagannathan and collaborators based on frequency and sleep grapho elements to detect EEG micro variations in alertness and characterize awake and drowsy trials.^76^ However, our PRL task design, especially the pretrial duration, which was limited to 2 seconds, did not fit the task features recommended by Jagannathan and collaborators (e.g., 4 seconds pretrial duration) for a reliable characterization of awake and drowsy trials. So we decided to classify awake/drowsy trials based on theta:alpha ratio, as seen in previous experiments from our laboratory.^17,30,71^

ii Deviation from pre-registration. Spearman rank-order correlation tests and Bayes factors were finally not performed to estimate the degree of association between switch likelihood as a function of consecutive negative feedbacks and arousal states. We will check the slope and effect.

iii Deviation from pre-registration. The mediation analysis was not initially included in the pre-registration, however, we decided to run it in order to test whether the impairment in performance in low and high arousal states could be attributed to the different maladaptive behavioural patterns.

## Notes

https://osf.io/xk379/

### Competing Interest Statement

The authors have declared no competing interest.

## REFERENCES

1. Salehi, B., Cordero, M. I. & Sandi, C. Learning under stress: The inverted-U-shape function revisited. Learn. Mem. 17, 522–530 (2010).

2. Yerkes, R. M. & Dodson, J. D. The relation of strength of stimulus to rapidity of habit-formation. J. Comp. Neurol. Psychol. 18, 459–482 (1908).

3. Shiu, L. & Chan, T. Unlearning a stimulus–response association. Psychol. Res. 70, 193–199 (2006).

4. Stalnaker, T. A., Cooch, N. K. & Schoenbaum, G. What the orbitofrontal cortex does not do. Nat. Neurosci. 18, 620 (2015).

5. Costa, V. D., Tran, V. L., Turchi, J. & Averbeck, B. B. Reversal Learning and Dopamine: A Bayesian Perspective. J. Neurosci. 35, 2407–2416 (2015).

6. Jang, A. I. et al. The role of frontal cortical and medial-temporal lobe brain areas in learning a Bayesian prior belief on reversals. J. Neurosci. 35, 11751–11760 (2015).

7. Izquierdo, A., Brigman, J. L., Radke, A. K., Rudebeck, P. H. & Holmes, A. The neural basis of reversal learning: An updated perspective. Neuroscience 345, 12–26 (2017).

8. Murray, E. A. & Gaffan, D. Prospective memory in the formation of learning sets by rhesus monkeys (Macaca mulatta). J. Exp. Psychol. Anim. Behav. Process. 32, 87 (2006).

9. Ciria, L. F. et al. Probabilistic trend detection in different levels of arousal. OSF (2019).

10. Goupil, L. & Bekinschtein, T. A. Cognitive processing during the transition to sleep. 15.

11. Schmit, C. & Brisswalter, J. Executive functioning during prolonged exercise: a fatigue-based neurocognitive perspective. Int. Rev. Sport Exerc. Psychol. 13, 21–39 (2020).

12. Ogilvie, R. D. & Wilkinson, R. T. The detection of sleep onset: behavioral and physiological convergence. Psychophysiology 21, 510–520 (1984).

13. Bernaola-Galván, P. A., Gómez-Extremera, M., Romance, A. R. & Carpena, P. Correlations in magnitude series to assess nonlinearities: Application to multifractal models and heartbeat fluctuations. Phys. Rev. E 96, 032218 (2017).

14. Wickens, C. D., Hutchins, S. D., Laux, L. & Sebok, A. The impact of sleep disruption on complex cognitive tasks: a meta-analysis. Hum. Factors 57, 930–946 (2015).

15. McMorris, T. & Hale, B. J. Differential effects of differing intensities of acute exercise on speed and accuracy of cognition: A meta-analytical investigation. Brain Cogn. 80, 338–351 (2012).

16. McMorris, T., Sproule, J., Turner, A. & Hale, B. J. Acute, intermediate intensity exercise, and speed and accuracy in working memory tasks: a meta-analytical comparison of effects. Physiol. Behav. 102, 421–428 (2011).

17. Noreika, V. et al. Alertness fluctuations when performing a task modulate cortical evoked responses to transcranial magnetic stimulation. NeuroImage 223, 117305 (2020).

18. Canales-Johnson, A. et al. Decreased alertness reconfigures cognitive control networks. J. Neurosci. 40, 7142–7154 (2020).

19. Nir, Y., Massimini, M., Boly, M. & Tononi, G. Sleep and consciousness. in Neuroimaging of consciousness 133–182 (Springer, 2013).

20. Cimenser, A. et al. Tracking brain states under general anesthesia by using global coherence analysis. Proc. Natl. Acad. Sci. 108, 8832–8837 (2011).

21. Hudson, A. N., Van Dongen, H. P. & Honn, K. A. Sleep deprivation, vigilant attention, and brain function: a review. Neuropsychopharmacology 45, 21–30 (2020).

22. Wise, R. A. Dopamine, learning and motivation. Nat. Rev. Neurosci. 5, 483–494 (2004).

23. Tanaka, M., Ishii, A. & Watanabe, Y. Neural correlates of central inhibition during physical fatigue. PloS One 8, e70949 (2013).

24. Chennu, S., O’Connor, S., Adapa, R., Menon, D. K. & Bekinschtein, T. A. Brain connectivity dissociates responsiveness from drug exposure during propofol-induced transitions of consciousness. PLoS Comput. Biol. 12, e1004669 (2016).

25. Lee, M. et al. Network properties in transitions of consciousness during propofol-induced sedation. Sci. Rep. 7, 1–13 (2017).

26. Pace-Schott, E. F. & Hobson, J. A. The neurobiology of sleep: genetics, cellular physiology and subcortical networks. Nat. Rev. Neurosci. 3, 591–605 (2002).

27. Bareham, C. A., Bekinschtein, T. A., Scott, S. K. & Manly, T. Does left-handedness confer resistance to spatial bias? Sci. Rep. 5, (2015).

28. Overgaard, M. & Overgaard, R. Measurements of consciousness in the vegetative state. The Lancet 378, 2052–2054 (2011).

29. Sanders, R. D., Tononi, G., Laureys, S. & Sleigh, J. Unconsciousness, not equal to unresponsiveness. Anesthesiology 116, 946–959 (2013).

30. Bareham, C. A., Manly, T., Pustovaya, O. V., Scott, S. K. & Bekinschtein, T. A. Losing the left side of the world: rightward shift in human spatial attention with sleep onset. Sci. Rep. 4, 1–5 (2014).

31. Jagannathan, S. R., Bareham, C. A. & Bekinschtein, T. A. Decreasing arousal modulates perceptual decision-making. http://biorxiv.org/lookup/doi/10.1101/2020.07.23.218727 (2020) doi:10.1101/2020.07.23.218727.

32. Chennu, S. & Bekinschtein, T. A. Arousal Modulates Auditory Attention and Awareness: Insights from Sleep, Sedation, and Disorders of Consciousness. Front. Psychol. 3, (2012).

33. Kouider, S., Andrillon, T., Barbosa, L. S., Goupil, L. & Bekinschtein, T. A. Inducing Task-Relevant Responses to Speech in the Sleeping Brain. Curr. Biol. 24, 2208–2214 (2014).

34. Berridge, C. W. & Arnsten, A. F. Psychostimulants and motivated behavior: arousal and cognition. Neurosci. Biobehav. Rev. 37, 1976–1984 (2013).

35. Kenney, W. L., Wilmore, J. H. & Costill, D. L. Physiology of sport and exercise. (Human kinetics, 2018).

36. Verburgh, L., Konigs, M., Scherder, E. J. A. & Oosterlaan, J. Physical exercise and executive functions in preadolescent children, adolescents and young adults: a meta-analysis. Br. J. Sports Med. 48, 973–979 (2014).

37. Chang, Y. K., Labban, J. D., Gapin, J. I. & Etnier, J. L. The effects of acute exercise on cognitive performance: A meta-analysis. Brain Res. 1453, 87–101 (2012).

38. Lambourne, K. & Tomporowski, P. The effect of exercise-induced arousal on cognitive task performance: A meta-regression analysis. Brain Res. 1341, 12–24 (2010).

39. Erickson, K. I., Hillman, C. H. & Kramer, A. F. Physical activity, brain, and cognition. Curr. Opin. Behav. Sci. 4, 27–32 (2015).

40. González-Fernández, F., Etnier, J. L., Zabala, M. & Sanabria, D. Vigilance performance during acute exercise. Int. J. Sport Psychol. 48, 435–447 (2017).

41. Andrillon, T., Poulsen, A. T., Hansen, L. K., Léger, D. & Kouider, S. Neural markers of responsiveness to the environment in human sleep. J. Neurosci. 36, 6583–6596 (2016).

42. Chmura, J., Krysztofiak, H., Ziemba, A. W., Nazar, K. & Kaciuba-Uścilko, H. Psychomotor performance during prolonged exercise above and below the blood lactate threshold. Eur. J. Appl. Physiol. 77, 77–80 (1997).

43. Del Giorno, J. M., Hall, E. E., O’Leary, K. C., Bixby, W. R. & Miller, P. C. Cognitive function during acute exercise: a test of the transient hypofrontality theory. J. Sport Exerc. Psychol. 32, 312–323 (2010).

44. Maran, T. et al. Lost in Time and Space: States of High Arousal Disrupt Implicit Acquisition of Spatial and Sequential Context Information. Front. Behav. Neurosci. 11, (2017).

45. Packard, M. G. & Goodman, J. Emotional arousal and multiple memory systems in the mammalian brain. Front. Behav. Neurosci. 6, 14 (2012).

46. Schwabe, L. & Wolf, O. T. Stress and multiple memory systems: from ‘thinking’to ‘doing’. Trends Cogn. Sci. 17, 60–68 (2013).

47. Broadhurst, P. L. The interaction of task difficulty and motivation: The Yerkes Dodson law revived. Acta Psychol. Amst. (1959).

48. Brown, J. S. Motivation of behavior. (Prabhat Prakashan, 1961).

49. Izquierdo, A. & Jentsch, J. D. Reversal learning as a measure of impulsive and compulsive behavior in addictions. Psychopharmacology (Berl.) 219, 607–620 (2012).

50. Swainson, R. et al. Probabilistic learning and reversal deficits in patients with Parkinson’s disease or frontal or temporal lobe lesions: possible adverse effects of dopaminergic medication. Neuropsychologia 38, 596–612 (2000).

51. Remijnse, P., Nielen, M., Uylings, H. & Veltman, D. Neural correlates of a reversal learning task with an affectively neutral baseline: An event-related fMRI study. NeuroImage 26, 609–618 (2005).

52. Clark, L., Cools, R. & Robbins, T. W. The neuropsychology of ventral prefrontal cortex: Decision-making and reversal learning. Brain Cogn. 55, 41–53 (2004).

53. Ersche, K. D., Roiser, J. P., Robbins, T. W. & Sahakian, B. J. Chronic cocaine but not chronic amphetamine use is associated with perseverative responding in humans. Psychopharmacology (Berl.) 197, 421–431 (2008).

54. Leeson, V. C. et al. Discrimination learning, reversal, and set-shifting in first-episode schizophrenia: stability over six years and specific associations with medication type and disorganization syndrome. Biol. Psychiatry 66, 586–593 (2009).

55. Kanen, J. W. et al. Effect of lysergic acid diethylamide (LSD) on reinforcement learning in humans. bioRxiv 2020–12 (2021).

56. den Ouden, H. E. M. et al. Dissociable Effects of Dopamine and Serotonin on Reversal Learning. Neuron 80, 1090–1100 (2013).

57. Murphy, F. C., Michael, A., Robbins, T. W. & Sahakian, B. J. Neuropsychological impairment in patients with major depressive disorder: the effects of feedback on task performance. Psychol. Med. 33, 455 (2003).

58. Dombrovski, A. Y., Szanto, K., Clark, L., Reynolds, C. F. & Siegle, G. J. Reward signals, attempted suicide, and impulsivity in late-life depression. JAMA Psychiatry 70, 1020–1030 (2013).

59. Dombrovski, A. Y. et al. Corticostriatothalamic reward prediction error signals and executive control in late-life depression. Psychol. Med. 45, 1413 (2015).

60. Tavares, J. V. T. et al. Neural basis of abnormal response to negative feedback in unmedicated mood disorders. Neuroimage 42, 1118–1126 (2008).

61. Mueller, E. M., Pechtel, P., Cohen, A. L., Douglas, S. R. & Pizzagalli, D. A. POTENTIATED PROCESSING OF NEGATIVE FEEDBACK IN DEPRESSION IS ATTENUATED BY ANHEDONIA: Research Article: Anhedonia and Feedback Processing in MDD. Depress. Anxiety 32, 296–305 (2015).

62. Drummond, S. The Effects of Total Sleep Deprivation on Cerebral Responses to Cognitive Performance. Neuropsychopharmacology 25, S68–S73 (2001).

63. Jackson, M. L. et al. Deconstructing and reconstructing cognitive performance in sleep deprivation. Sleep Med. Rev. 17, 215–225 (2013).

64. Gevers, W., Deliens, G., Hoffmann, S., Notebaert, W. & Peigneux, P. Sleep deprivation selectively disrupts top-down adaptation to cognitive conflict in the Stroop test. J. Sleep Res. 24, 666–672 (2015).

65. Whitney, P. et al. Sleep deprivation diminishes attentional control effectiveness and impairs flexible adaptation to changing conditions. Sci. Rep. 7, 1–9 (2017).

66. Whitney, P., Hinson, J. M. & Nusbaum, A. T. A dynamic attentional control framework for understanding sleep deprivation effects on cognition. Prog. Brain Res. 246, 111–126 (2019).

67. Bekinschtein, T. A. et al. Neural signature of the conscious processing of auditory regularities. Proc. Natl. Acad. Sci. 106, 1672–1677 (2009).

68. Teigen, K. H. Yerkes-Dodson: A Law for all Seasons. Theory Psychol. 4, 525–547 (1994).

69. Mauss, I. B. & Robinson, M. D. Measures of emotion: A review. Cogn. Emot. 23, 209–237 (2009).

70. Tagliazucchi, E. Decoding Wakefulness Levels from Typical fMRI Resting-State Data Reveals Reliable Drifts between Wakefulness and Sleep. 14.

71. Comsa, I. M., Bekinschtein, T. A. & Chennu, S. Transient Topographical Dynamics of the Electroencephalogram Predict Brain Connectivity and Behavioural Responsiveness During Drowsiness. Brain Topogr. 32, 315–331 (2019).

72. Johns, M. W. A new method for measuring daytime sleepiness: the Epworth sleepiness scale. sleep 14, 540–545 (1991).

73. Holgado, D. et al. Tramadol effects on physical performance and sustained attention during a 20-min indoor cycling time-trial: A randomised controlled trial. J. Sci. Med. Sport 21, 654–660 (2018).

74. Holgado, D. et al. Transcranial direct current stimulation (tDCS) over the left prefrontal cortex does not affect time-trial self-paced cycling performance: Evidence from oscillatory brain activity and power output. PloS One 14, e0210873 (2019).

75. Zandonai, T. et al. Novel evidence on the effect of tramadol on self-paced high-intensity cycling. J. Sports Sci. 1–9 (2021) doi:10.1080/02640414.2021.1877440.

76. Jagannathan, S. R. et al. Tracking wakefulness as it fades: Micro-measures of alertness. NeuroImage 176, 138–151 (2018).

77. Saper, C. B., Fuller, P. M., Pedersen, N. P., Lu, J. & Scammell, T. E. Sleep State Switching. Neuron 68, 1023–1042 (2010).

78. Ciria, L. F., Perakakis, P., Luque-Casado, A. & Sanabria, D. Physical exercise increases overall brain oscillatory activity but does not influence inhibitory control in young adults. NeuroImage 181, 203–210 (2018).

79. Ciria, L. F. et al. Oscillatory brain activity during acute exercise: Tonic and transient neural response to an oddball task. Psychophysiology e13326 (2019) doi:10.1111/psyp.13326.

80. Tanaka, H., Monahan, K. D. & Seals, D. R. Age-predicted maximal heart rate revisited. J. Am. Coll. Cardiol. 37, 153–156 (2001).

81. Bates, D. et al. Package ‘lme4’. Convergence 12, (2015).

82. Tingley, D., Yamamoto, T., Hirose, K., Keele, L. & Imai, K. mediation : *R* Package for Causal Mediation Analysis. J. Stat. Softw. 59, (2014).

